# Translational control of microglial inflammatory and neurodegenerative responses

**DOI:** 10.1101/2024.04.06.587750

**Authors:** Sara Bermudez, Jung-Hyun Choi, Jacob W. Vogel, Sung-Hoon Kim, Niaz Mahmood, Vivian Yuchan Zhu, Danielle Cozachenco, Moein Yaqubi, Linqiao Zhou, Jo Ann Stratton, Oskar Hansson, Luke Healy, Argel Aguilar-Valles, Nahum Sonenberg

## Abstract

In Alzheimer’s Disease (AD), activation of the mechanistic target of rapamycin (mTOR) pathway is essential for microglia neuroprotective roles, but it is unclear which mTOR effectors promote these neuroprotective functions. The mTOR complex 1 (mTORC1) inactivates the translation suppressors eukaryotic translation Initiation Factor 4E (eIF4E)-Binding Proteins (4E-BP) to promote mRNA translation. We show that 4E-BP1 inactivation is impaired in microglia under AD-relevant conditions. Depleting 4E-BPs in microglia increases mitochondrial metabolism, suppresses the pro-inflammatory profile, and mitigates amyloid-induced apoptosis. Furthermore, in the cerebrospinal fluid of patients with amyloid pathology, there was a positive association between microglia activation and neurodegeneration, which increases along 4E-BP1 levels. Thus, we propose the engagement mTORC1-4E-BP1 axis as a neuroprotective mechanism and a therapeutic target or biomarker for microglia modulation in AD.

---

Pathologic amyloid-β (Aβ) accumulation precedes cognitive decline in AD (1, 2). Microglia play a neuroprotective role via receptors that bind pathological forms of Aβ and transduce intracellular signals to induce phagocytosis and compaction of larger Aβ deposits into inert plaques (3–5). AD-risk genes are enriched in microglia receptors and signal transducers, including Triggering Receptor Expressed on Myeloid Cells 2 (TREM2), thus underscoring the linkage between AD genetic susceptibility and microglia function (6–8). During the AD disease course, homeostatic microglia react to pathology and acquire different states, such as disease-associated microglia (DAM), exhibiting unique transcriptional signatures (9, 10). However, some microglia populations lose their intrinsic protective functions during AD, possibly due to chronic inflammatory stimulation by Aβ aggregates or failure of physiological immune resolution mechanisms (11, 12). Indeed, in post-mortem brains, microglia adjacent to Aβ plaques exhibit dystrophic and fragmented morphology, consistent with the notion that AD progression is associated with reduced neuroprotective microglial function (13). Although ample evidence supports a role for both the neuroprotective role of microglia in AD and its ultimate loss, microglia functions depend on complex intracellular signaling pathways and the precise molecular pathways that underly these processes are not completely understood (9, 14–16). Identifying the molecular pathways that govern these functions is necessary to uncover avenues to boost microglia neuroprotective roles to ameliorate AD.

Activation of the mTOR pathway is pivotal to microglial survival and neuroprotective functions (17–20). Indeed, mTOR signaling is impaired in microglia deficient in TREM2 function or in the presence of amyloidosis (17, 18). Furthermore, a mTORC1-related metabolic deficit (e.g., energetic deficit, increased autophagy, unresponsiveness to stimuli, and apoptosis) is associated with microglia dysfunction in AD (21). A primary downstream target of the mTORC1 pathway is mRNA translation. mTORC1 induces phosphorylation and inactivation of eukaryotic initiation factor 4E (eIF4E)-binding proteins (4E-BPs), allowing their dissociation from elF4E and, thus, promoting translation (22). 4E-BPs preferentially suppress the translation of a subset of mRNAs, including those encoding mitochondrial proteins, thus dramatically inhibiting cellular metabolism (23). Of the three 4E-BP isoforms, 4E-BP1 is the most prominent in microglia, followed by 4E-BP2, while there is negligible expression of 4E-BP3 (24–26). 4E-BP1 has been proposed as a disease-specific biomarker in AD (27). Notably, inflammatory stimuli activate 4E-BP1 transcription in microglia in vivo (24, 28). Consistent with this, microglial activation markers and disease progression in AD strongly correlate with increased 4E-BP1 levels in cerebrospinal fluid in AD patients (29, 30). Although this association suggests a role of 4E-BP1 in the loss of the protective function of microglia, the mechanisms underlying this association remain to be elucidated. To address this gap, we investigated the mechanisms at the intersection of mTORC1-4E-BP-regulated mRNA translation and metabolism in the microglial response to Aβ. We demonstrate that dysregulation of 4E-BPs in microglia drives a dysfunctional and detrimental state that could lead to increased neurodegeneration. Thus, mTORC1 signaling critically impacts microglia physiology and promotes neuroprotective functions via 4E-BP-dependent mRNA translation.

## TREM2/SYK engages the mTORC1-4E-BP1 pathway

TREM2 signaling and microglia functions have been extensively documented in the BV2 microglia cell line (31–33). We first examined the effects of exposure to Aβ oligomers (Aβo) on BV2 microglia.

Acute exposure to Aβo (2 μM, 45 min) triggered phosphorylation of 4E-BP1 in microglia (Fig. 1A). Prolonged Aβo exposure (2 μM, 24h) was accompanied by an inflammatory response, as measured by increased expression of pro-IL-1β (Fig. S1). Concomitantly, there was an increase in the total levels of 4E-BP1 following a 24h Aβo exposure.

**Fig. 1.**
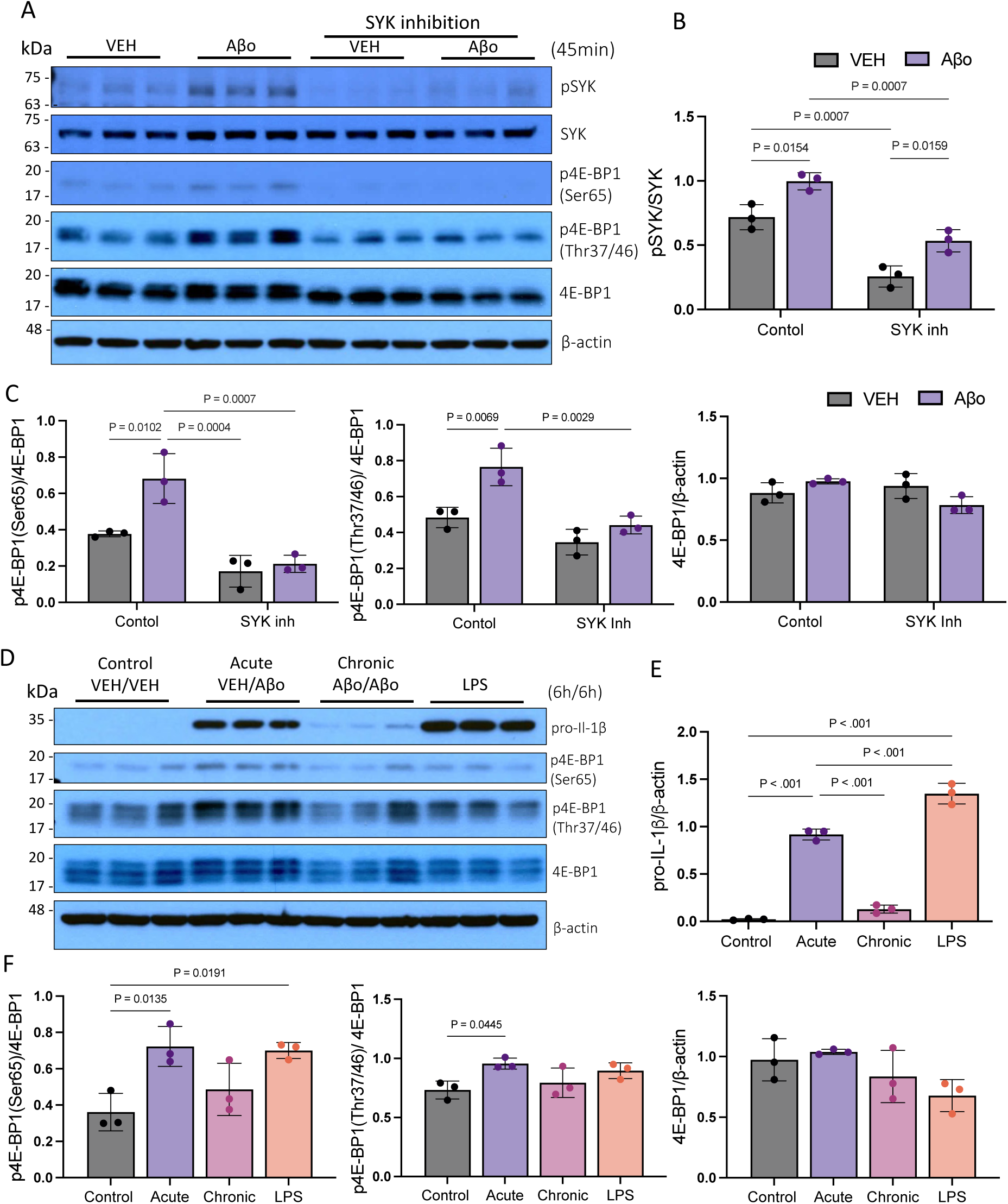
Acute AβO exposure triggers 4E-BP1 phosphorylation, whereas it is decreased by SYK signaling inhibition or chronic Aβo exposure. (A) Representative western blot of total and phosphorylated SYK and 4E-BP1 in BV2 cell lysates 45 minutes after vehicle (control) or Aβo (2 μM) treatment with or without SYK inhibitor (1 μM, R406). (B and C), Quantification of phosphorylation ratio to total protein or to β-actin loading control (n=3/group). (D) Representative western blot analysis of pro-IL-1β and total and phosphorylation status of 4E-BP1 in cell lysates from BV2 microglia after treatment with vehicle (control), acute, chronic (repeated) Aβo (2μM) or LPS (100ng/μL). (E) Quantification of pro-IL-1β levels and (F) phosphorylation ratio to total protein or to β-actin loading control (n=3/group). Data are presented as means ± SEM. (P values were obtained using two-way ANOVA, Tukey’s post-hoc test (B and C), and one-way ANOVA, Tukey’s post-hoc test (E and F). Samples in each group are from independent wells examined in the same blot.

Aβos bind to TREM2 on microglia with high affinity and elicit a robust inflammatory response (17, 34–37). To examine the role of the TREM2 pathway in the acute Aβo-induced phosphorylation of 4E-BP1, microglia were exposed to Aβo in the presence or absence of a Spleen Tyrosine Kinase (SYK) inhibitor (R406). Once recruited to DNAX Activating Protein of 12 kDa (DAP12), downstream of TREM2, SYK is phosphorylated (38). SYK depletion or inhibition reduces mTOR signaling and microglia’s neuroprotection in vitro and in vivo (20, 39, 40). As expected, exposure to Aβo elicited a rapid ∼40% increase in SYK phosphorylation (Fig 1B). Baseline and Aβo-induced SYK phosphorylation were reduced by ∼2-fold by a SYK inhibitor (Fig. 1B). Inhibition of SYK led to a ∼2-fold decrease in 4E-BP1 phosphorylation at serine 65, and the priming sites, threonines 37/46 (Fig. 1C). Phosphorylation at serine 65 is dependent on the phosphorylation of threonines 37/46, and is necessary for 4E-BP1 release of eIF4E (41). Thus, activation of TREM2/SYK by Aβo stimulates mTORC1-dependent translation.

Chronic Aβo exposure causes microglia dysfunction, reduced mTOR activity, and deficient energetic metabolism (17, 42). To determine the impact of chronic Aβo exposure on the mTORC1-4E-BP axis signaling, microglia were repeatedly exposed to Aβo (2 μM, 2x 6h, Extended Data Fig. 2) (17). A single exposure to Aβo or LPS (100 ng/ml, 1x 6h) induced a 2-fold increase in 4E-BP1 phosphorylation of Serine 65 (Fig. 1D) and an inflammatory response as reflected in elevated expression of pro-IL-1β (Fig. 1E). In contrast, in cells repeatedly treated with Aβo (2 μM, 2x 6h), pro-IL-1β was no longer induced, indicating immune tolerance, which occurred alongside a lack of induction in 4E-BP1 phosphorylation (Fig. 1F). The results indicate occlusion of the mTORC1-4E-BP axis in response to repeated Aβo exposure, and, consequently, deficient induction of mRNA translation and cellular metabolism. This effect could be associated with the microglia dysfunctional phenotype observed during amyloidosis. Consequently, it is conceivable that alleviating 4E-BP-mediated translation suppression should rescue microglial metabolic capacity and neuroprotective functions.

## Depletion of 4E-BPs in microglia increases mitochondria biogenesis and decreases inflammatory output

The mTORC1-4E-BPs signaling axis controls mitochondrial activity and biogenesis by selectively promoting the translation of cytoplasmic mRNAs encoding for mitochondrial proteins (23). Importantly, TREM2-deficient microglia exhibit reduced mitochondrial mass (18). Thus, we examined the metabolic profile of BV2 microglial cells lacking 4E-BP1 and 4E-BP2. We generated 4E-BP1 and 4E-BP2 double knockout (DKO) in BV2 microglia using CRISPR-CAS9 (Extended data Fig. 3). Mitochondrial mass in DKO microglia was increased by ∼50% compared to WT as determined by mitochondrial staining followed by confocal imaging (Fig. 2A). Next, we assessed mitochondrial respiratory function by real-time measurement of oxygen consumption rate (OCR). The maximal respiration capacity of the DKO cells was increased by 33% compared to WT (Fig. 2B).

**Fig. 2.**
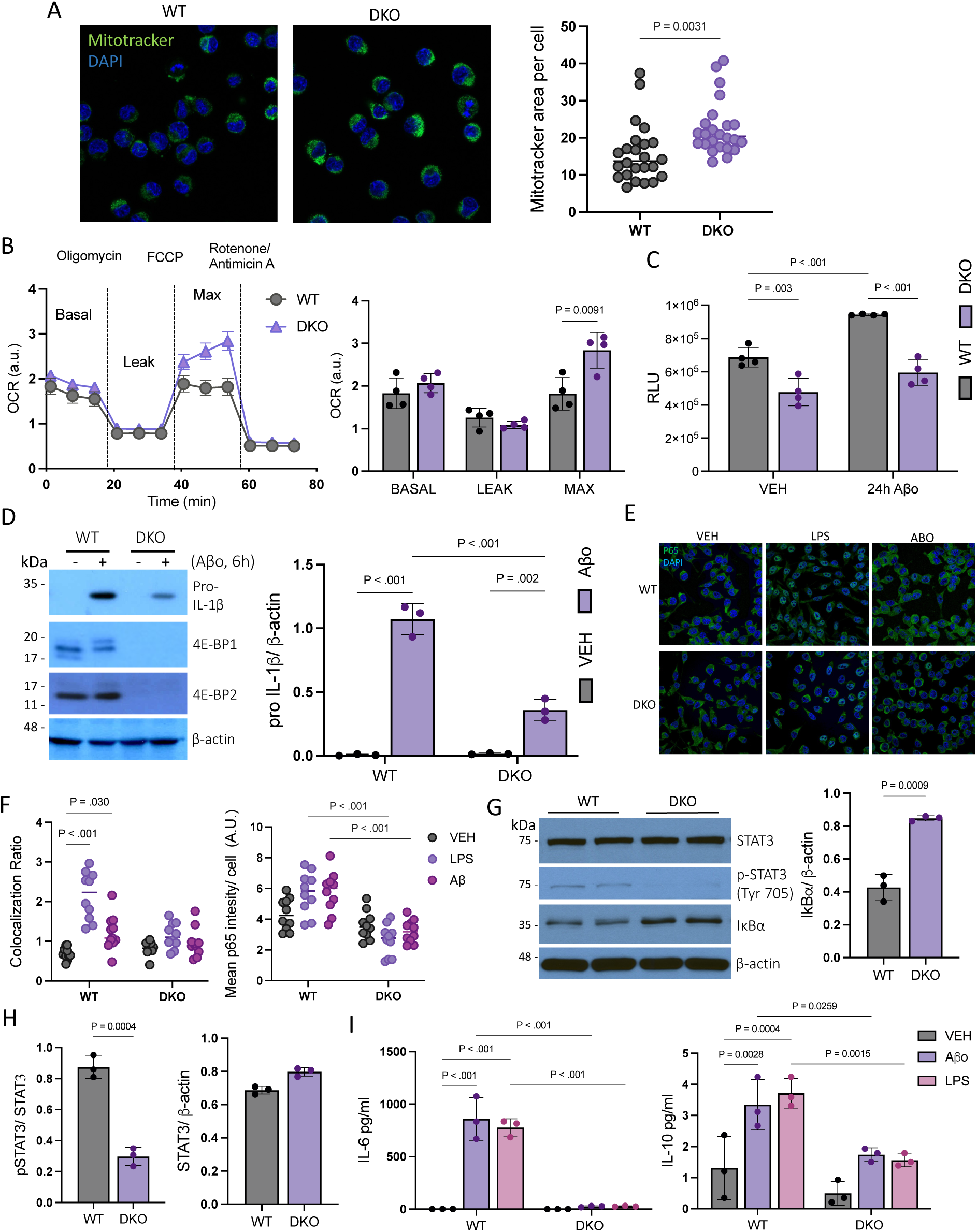
4E-BPs depletion in microglia increases mitochondrial mass, promotes oxidative phosphorylation, and suppresses pro-inflammatory responses. (A) Representative confocal images of microglia mitochondria labeled with MitoTracker™ and quantification of green fluorescence area (n=24, from 3 independent wells/group). (B) Real-time measurement of oxidative phosphorylation measured by oxygen consumption rate (OCR) (n=4 independent wells/group). a.u; arbitrary units. (C) Glycolysis level determined by lactate secretion 24h after Aβo (2μM) or vehicle control treatment (n=4 independent wells/group). RLU; relative light units. (D) Western blot analysis and quantification of pro-Il-1β 6h after Aβo exposure, and 4E-BP1 and 4E-BP2 Knock out confirmation (n=3, from 3 independent wells/group). (E) Representative confocal microscopy images of p65 cellular localization and fluorescence intensity, and (F) quantification of nuclear co-localization and total fluorescence intensity after vehicle control, Aβo (24h) or LPS (6h) stimulation (n=10, from 3 independent wells/group). (G) Representative western blot of WT and DKO microglia for STAT3, IκBα, and (H) quantification phosphorylated and total STAT3. (I) Determination of cytokines in supernatant from Aβo (24h) or LPS (6h) treated microglia. The colocalization ratio represents the total cytoplasmic p65 fluorescence intensity over DAPI. Data are presented as means ± SEM (P values were calculated using two-tailed unpaired t-test (A, G and H), two-way ANOVA, Tukey’s posthoc test (B, C, D, F and I))

**Fig. 3.**
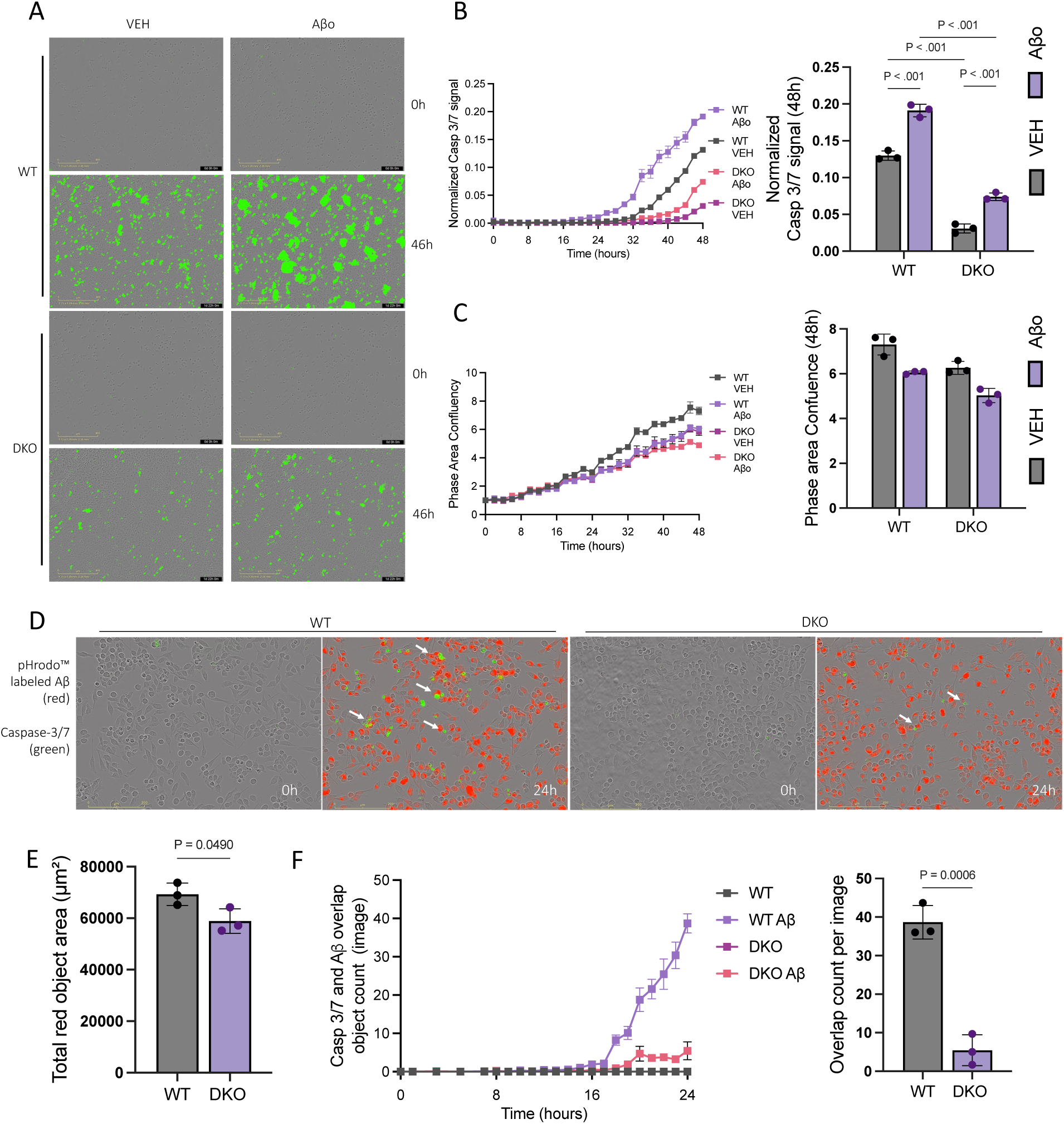
Depletion of 4E-BPs mitigates microglia apoptosis associated with Aβ. (A) Representative IncuCyte micrographs of WT and DKO microglia co-incubated with Aβo or vehicle control and CellEvent™ Casp3/7 (green) at 0 and 46h. (B) Measurement of Casp3/7 green fluorescence over phase area for 48h and quantification at 48h (n=3 independent wells/group with 4 images per well). (C) Confluence measurement for 48h and quantification at 48h (n=3 independent wells/group with 4 images per well). (D) Representative IncuCyte micrographs of microglia co-incubated with pHrodo^™^ labeled Aβ or vehicle (control). (E) quantification of total red fluorescence area per image at 24h (n=3 independent wells/group with 4 images per well). (F) quantification of Casp3/7 (green) and pHrodo^™^ labeled Aβ (red) co-labeled cells over 24h and quantification at 24h. Data are presented as means ±SEM. (P values were calculated using two-way ANOVA, Tukey’s post-hoc test (B and C), and unpaired t-test (E and F)).

In microglia, like in macrophages, immune activation is linked to metabolic reprogramming from oxidative phosphorylation (OXPHOS) to glycolysis (43, 44). To investigate how 4E-BPs’ depletion changes the glycolytic switch, we measured lactate secretion as an indicator of glycolysis in BV2 cells. DKO microglia show a 30% reduction in lactate secretion, indicative of diminished reliance on glycolytic metabolism in the DKO cells compared to WT (Fig. 2C). Aβo treatment (2 μM, 24h) caused a 38% increase in glycolysis in WT but not in DKO microglia. Given the role of metabolism modulating immune responses in microglia, we next examined the impact of 4E-BPs deletion on the inflammatory profile of DKO BV2 cells in response to Aβo treatment. Aβo treatment (2 μM, 6h) resulted in a 67% lower induction of pro-IL-1β in the DKO microglia compared to WT (Fig. 2D). Next, we investigated the status of major transcription factors associated with the inflammatory response in microglia such as signal transducer and activator of transcription 3 (STAT3) and nuclear factor kappa B (NFκB), which induces pro-IL-1β expression (45). Immunocytochemistry was performed to determine the cellular localization of the NFκB subunit p65 in WT and DKO cells stimulated with Aβo (2 μM, 24h) or LPS (100ng/ml, Fig. 2e). Translocation of p65 into the nucleus was observed in the WT but not in the DKO cells after stimulation with either Aβ or LPS (Fig. 2F). Furthermore, p65 amount was decreased in DKO compared to WT upon Aβ or LPS treatment (by 44% and 54%, respectively). This decrease was compounded by a 2-fold increase in the NFκB inhibitor, IkBα, in DKO cells (Fig. 2G). In addition, at baseline, STAT3 phosphorylation was decreased in DKO cells by 66% (Fig. 2H). Overall, 4E-BPs depletion reduced the activity of transcription factors associated with the microglia inflammatory response.

To further demonstrate that the inflammatory response was attenuated in DKO cells, we measured cytokine levels released from BV2 cells treated with Aβo (2 μM, 24h) or LPS (100ng/ml, 6h). IL-6 and IL-10 were elevated in WT microglia but not in the DKO cells (Fig. 2I and Fig. S4). Although a significant induction of pro-IL-1β in WT microglia was observed, IL-1β was not detected in the culture media, perhaps because IL-1β secretion requires induction of inflammasome assembly by a second stimulus (46). Thus, consistent with the decreased glycolytic switch in DKO microglia (Fig. 2C), 4E-BPs depletion caused a reduction in the induction of several pro-inflammatory cytokines in microglia.

## 4E-BPs depletion mitigates microglial apoptosis

Microglia harboring TREM2 mutations exhibit metabolic deficit, and hypersensitivity to stress-induced apoptosis (18, 47, 48). Thus, we investigated the role of 4E-BPs in Aβo-induced apoptosis by comparing the apoptotic response in WT and DKO microglia. We exposed microglia to Aβo (2µM) and measured apoptosis by caspase 3/7 cleavage in live cells for 48h (Fig. 3A). Aβo induced apoptosis significantly more (61%) in WT than DKO microglia. The decrease in apoptosis was similarly observed even without Aβo stimulation, starting after 24 hours without media change (Fig. 3B), whereas proliferation was not affected by 4E-BP depletion (Fig. 3C).

Given the reduced glycolytic induction, inflammatory response, and apoptosis in DKO microglia exposed to Aβo, we examined whether DKO microglia could phagocytize Aβo. To this end, we exposed DKO and WT microglia to pHrodo-labeled Aβ (1µM) while simultaneously measuring apoptosis by caspase 3/7 cleavage in live cells for 24h to determine whether Aβo phagocytosis colocalizes with apoptotic cells (Fig. 3D). We found that both WT and DKO microglia phagocytized pHrodo-labeled Aβo (Fig. 3E). Co-localization with caspase 3/7 activity was decreased by ∼85% in DKO cells compared to WT (Fig. 3D and F). The results show that deletion of 4E-BPs mitigates microglia apoptosis associated with Aβo phagocytosis. This underscores the importance of engaging the mTOR-4E-BP1 axis for microglia survival in response to amyloidosis.

## The expression of various inflammatory mediators is reduced in 4E-BP-depleted microglia in vivo

We next examined the impact of 4E-BPs depletion on the microglial translatome in vivo. We used a Cre-dependent ‘Ribo-Tag’ mouse strain (Rpl22^HA/+^), allowing for immunoprecipitation of ribosome-bound mRNAs. The mouse strain was crossed with a microglia-specific Cre mouse model with or without LoxP sequences flanking the Eif4ebp1 and Eif4ebp2 genes (Microglia [MG]^DKO^: Cx3cr1^CreERT2/+^; Eif4ebp1 ^f/f^ Eif4ebp2 ^f/f^; Rpl22^HA/+^ or MG^WT^: Cx3cr1^CreERt2/+^; Rpl22^HA/+^) to perform microglia specific ribosome-bound mRNAs immunoprecipitation (49) (Fig. 4A and Fig. S5). After mRNA isolation, sequencing was performed. Cell-type deconvolution, using the Allen Mouse Brain “Whole cortex & hippocampus with 10x-smart-seq taxonomy” atlas as the reference dataset, validated the immunoprecipitation and microglia translatome enrichment ( Fig. S6)(50).

**Fig. 4.**
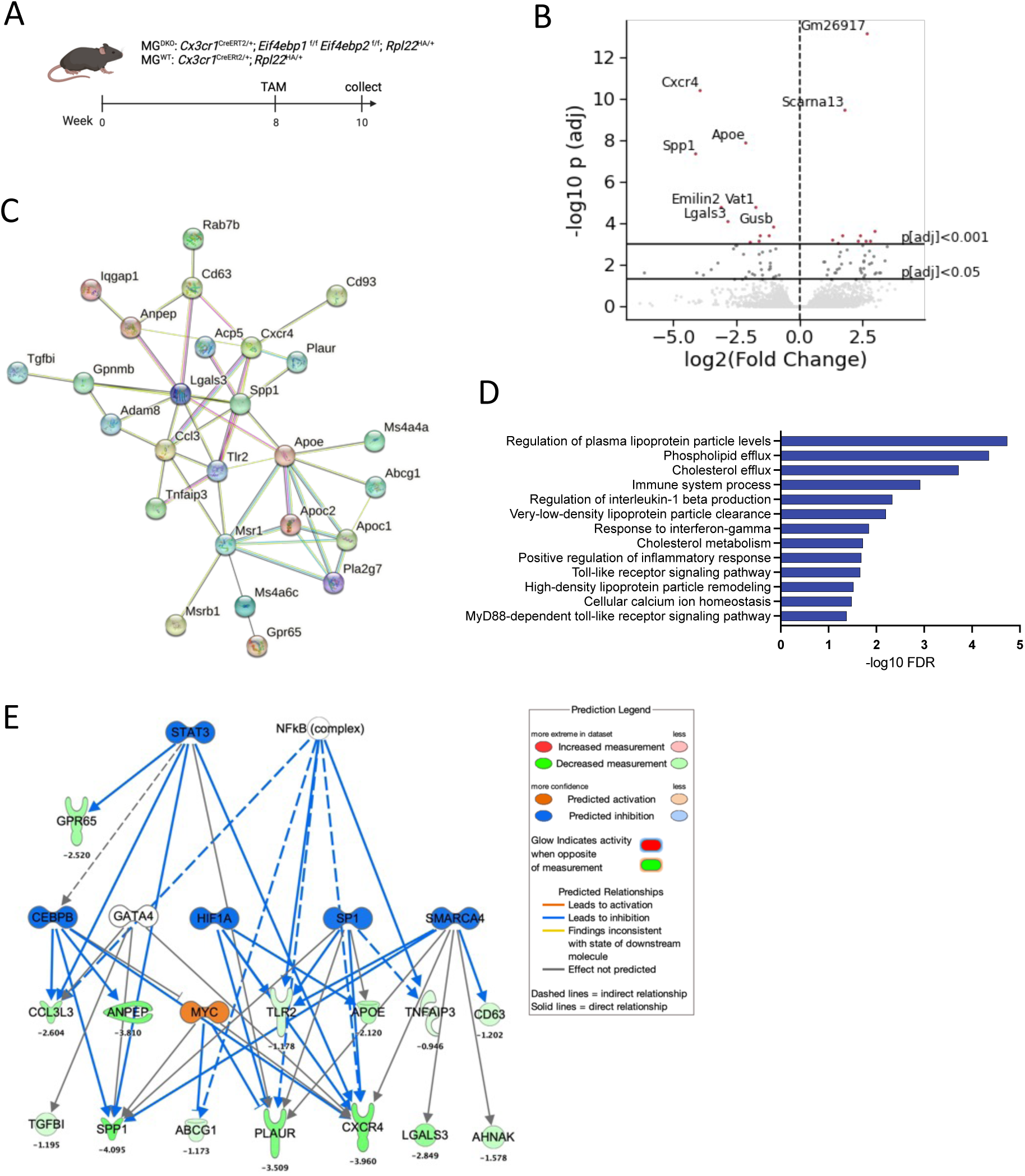
Microglia-specific depletion of 4E-BPs induces baseline changes in the microglia translatome in vivo. (A) Schematic diagram illustrating mouse crossing and tamoxifen (TAM) induction timeline. (B) Volcano plot of changes in microglia translatome upon 4E-BP depletion (n=2 to 3 independent samples/group). (C) String database protein-protein interaction network of downregulated DEGs in MG^DKO^ vs. MG^WT^ translatome. (D) String database functional enrichment analyses of downregulated DEGs in MG^DKO^ translatome. (E) Ingenuity pathway analysis (IPA) network of upstream regulators (transcription factors) generated using downregulated DEGs in MG^DKO^. We report genes surviving multiple comparisons correction (p[adj]<0.05).

Next, an analysis of the translatome of MG^DKO^ and MG^WT^ was carried out to identify differentially expressed genes (DEGs, p[adj]<0.05) (Fig. 4B). In agreement with the in vitro findings, the translatome from the MG^DKO^ mice showed downregulation of several mediators associated with microglia disease-associated phenotypes compared to the MG^WT^ translatome. These include genes encoding apolipoprotein E (Apoe), toll-like receptor 2 (Tlr2), CC-chemokine ligand 3 (Ccl3), and osteopontin (Spp1) (9, 28, 51). The STRING database predicted downregulated DEGs in MG^DKO^ to be a highly associated network (Protein-Protein Interaction enrichment p-value: < 1.0e-16) (Fig. 4C). STRING database functional enrichment analyses showed that the depletion of 4E-BPs in microglia in vivo engendered a decrease in several biological functions related to apolipoprotein production and regulation, as well as inflammation-associated functions such as IL-1β regulation and positive regulation of the inflammatory response. The downregulation is consistent with the decreased pro-inflammatory profile of microglia lacking 4E-BPs, even in the absence of Aβ stimulation (Fig. 4D). No significant functional enrichment was detected in upregulated DEGs. Ingenuity pathway analysis (IPA) of upstream regulators (transcription factors) was performed based on differentially expressed genes between MG^WT^ vs. MG^DKO^ (with corresponding fold changes, Fig. 4E). Consistent with the in vitro data, the upstream regulator analysis predicted the inhibition of STAT3 activity (activation z-score of −2.376) in the MG^DKO^ compared to MG^WT^. Although the NFκB complex was predicted to be inhibited, differences between strains did not reach a significant score (activation z-score of −1.64, an absolute z-score of ≥ 2 is considered significant).

## 4E-BP1 levels in CSF moderate the relationship between microglial activation and neurodegeneration

4E-BP1 has been proposed as a disease-specific biomarker in AD (27). 4E-BP1 levels in CSF of AD patients are positively correlated with microglia activation markers and worsened AD pathology (29, 30). Our in vitro results highlight the role of 4E-BPs inhibition in limiting the microglial pro-inflammatory responses. We sought to complement these findings using measurements of 4E-BP1 in CSF of healthy and AD-spectrum individuals to assess whether 4E-BP1 influenced the effect of microglial activation in neurodegeneration. We used soluble TREM2 (sTREM2) levels, a marker of microglia activation, and Neurofilament light (NfL), a widely used measure of neurodegeneration (52, 53). There was a significant interaction (β=6.00 [0.16,11.83], t= 2.02, p=0.044) between 4E-BP1 levels and soluble sTREM2 on NfL levels in CSF. Importantly, stronger positive associations between sTREM2 and NfL were observed in association with higher 4E-BP1 levels (Figure 5A). This interaction was driven by individuals harboring brain β-amyloid pathology (Fig. 5B). These data support the role of 4E-BP1 as a driver and potential biomarker of detrimental microglial states in response to β-amyloid pathology.

**Fig. 5.**
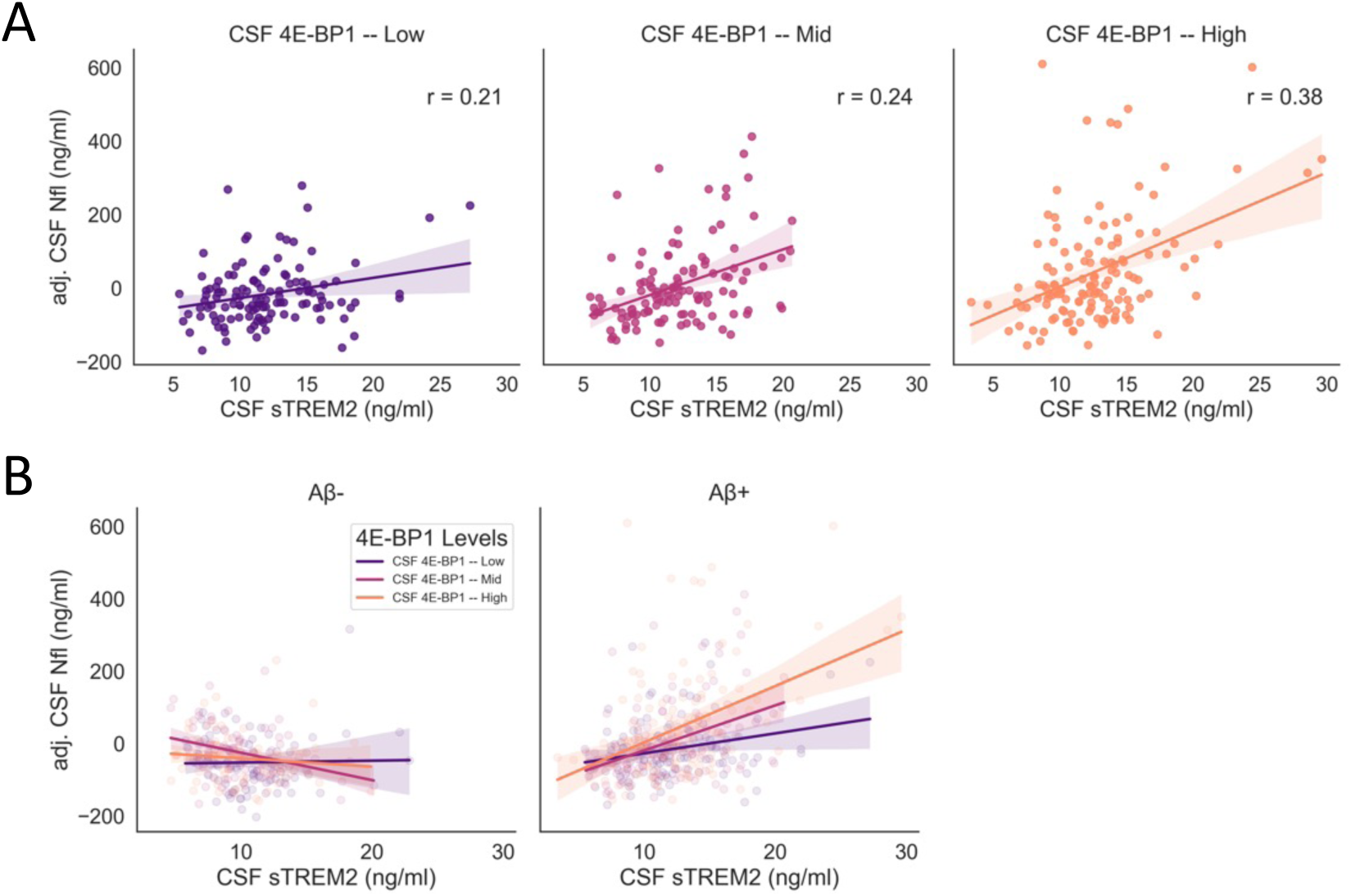
4E-BP1 levels in CSF moderate the association between microglia activation and neurodegeneration in AD patients with amyloid pathology. (A) Scatter plot representing concentrations in CSF of Nfl as a function of sTREM2 concentration in the different 4E-BP1 categories (either high, middle or low concentrations of 4E-BP1 in CSF). (B) Scatter plot representing concentrations in CSF of Nfl as a function of sTREM2 concentration in the different 4E-BP1 categories in individuals with or without Aβ pathology. Solid lines indicate the linear regression and values are indicated. r; correlation coefficient.

## Discussion

Here, we demonstrate that the mTORC1-4E-BP1 axis is a central effector of microglia physiology in AD-relevant conditions. First, in vitro experiments showed that Aβo trigger the mTORC1-mediated inactivation of 4E-BP1, which is dependent on SYK intracellular signaling and is decreased during chronic exposure to Aβo (Fig. 1). 4E-BP1 expression was induced during prolonged exposure, suggesting that the increased expression may surpass the mTORC1-mediated inactivation (Fig. S1). Deletion of 4E-BPs (DKO) stimulates mitochondrial biogenesis in microglia and increases mitochondrial respiration capacity (Fig. 2A and B) while decreasing glycolytic induction upon Aβo exposure (Fig. 2C). The suppressed glycolytic switch in DKO microglia was associated with decreased induction of pro-IL-1β and several inflammatory cytokines (Fig. 2D-I), and reduced Aβo-phagocytosis-induced apoptosis in 4E-BP-depleted microglia (Fig. 3). In vivo, microglial knockout of the 4E-BPs in mice induced a downregulation in select genes related to microglial disease-associated phenotypes (Fig. 4). Finally, we showed a stronger positive association of microglia activation with neurodegeneration in the presence of higher levels of 4E-BP1 in cerebrospinal fluid of patients with β-amyloid pathology (Fig. 5). Thus, we propose that the absence of mTORC1-mediated 4E-BP1 inactivation, or increased 4E-BP1 expression engenders dysfunctional or detrimental states in microglia, leading to increased neurodegeneration.

In our analysis of microglia in vivo translatome, 4E-BP depletion did not affect classic microglia markers or homeostatic signature gene expression. However, we noted a decrease in levels of Apoe and several genes linked to microglial disease-associated phenotypes in MG^DKO^ compared to MG^WT^. The effects may be an indirect consequence of the translational or post-translational regulation of key transcription factors or their regulators by the absence of 4E-BPs in microglia (54, 55). Apoe is one of the most abundantly translated mRNAs in microglia, even at basal level, and is increased in neurodegenerative diseases as a core component of DAM states (28). Thus, our findings indicate a modulatory role of the mTORC1-4E-BP1 axis on microglia disease-associated phenotype gene expression. In this regard, SYK-deficient microglia acquire an Apoe-expressing prodromal stage of DAM but fail to support this phenotype metabolically or perform neuroprotective functions (20). Considering the strong impact of SYK signaling on 4E-BP1 phosphorylation, we propose that when activated, the SYK-mTORC1-4E-BP axis sustains mitochondrial metabolism and attenuates the induction of disease-associated phenotypes. This mechanism allows the maintenance of cell viability and limits inflammatory responses while conserving microglia neuroprotective functions such as Aβ phagocytosis.

Mitochondrial metabolism is a key regulator of microglia phenotypic transitions, including the regulation of transcription factors associated with inflammation (44). For example, over-expression of mitochondrial proteins like TFAM (transcription factor A, mitochondrial) reduces NF-kB nuclear translocation and IL-1b expression in vivo (56). Consistent with the observed changes in mitochondrial metabolism following 4E-BP depletion in vitro, DKO microglia in vitro and in vivo exhibited a marked decrease in the baseline and inflammation-induced activity of transcription factors associated with inflammatory response, such as STAT3 and NFκB. Thus, mTOR profoundly affects microglia translatomic and transcriptomic landscape through 4E-BP-regulated mitochondrial metabolism.

We found a stronger positive association between microglia activation (sTREM2) and neurodegeneration (NfL) in the presence of higher levels of 4E-BP1 in the CSF, specifically in patients with brain β-amyloid pathology. TREM2 cleavage into sTREM2 can decrease downstream signaling transduction to mTOR activation and thus reduce 4E-BP1 inactivation (57), highlighting the potential involvement of 4E-BP1 in microglia inflammatory and neurodegenerative responses. In this regard, our pre-clinical findings indicate that mTORC1-mediated inactivation of 4E-BP1 in microglia plays a critical role in limiting microglia pro-inflammatory output by pivoting the associated glycolytic energy pathway towards OXPHOS-dependent metabolism, ultimately acting as a neuroprotective mechanism of immune resolution (Fig. S7). Taken together, our study underscores the importance of mRNA translation downstream of mTORC1 signaling in microglia and its implication in AD. As such, 4E-BP1 is an appealing target for therapy in AD, and its targeting to modulate microglia and monitoring is a potential novel avenue deserving of clinical investigation.

## Supporting information

DEGs DKO vs WT translatome

## Acknowledgments

We thank A. Sylvestre, A. Lafrance, I. Harvey and E. Migon for technical assistance.

## Funding

FRQNT (Fonds de recherche du Québec – Nature et technologies) doctoral fellowship (SB)

HBHL (Healthy Brains, Healthy Lives) doctoral fellowship (SB)

CIHR Foundation Grant FND-148423 (NS)

Alzheimer Society of Canada grant (NS)

ERA PerMed ERAPERMED2021-184 (OH)

The Knut and Alice Wallenberg foundation 2017-0383 (OH)

The Strategic Research Area MultiPark (Multidisciplinary Research in Parkinson’s disease) at Lund University (OH)

The Swedish Alzheimer Foundation AF-980907 (OH)

The Swedish Brain Foundation FO2021-0293 (OH)

The Parkinson foundation of Sweden 1412/22 (OH)

The Cure Alzheimer’s fund, the Konung Gustaf V:s och Drottning Victorias Frimurarestiftelse, the Skåne University Hospital Foundation 2020-O000028 (OH)

Regionalt Forskningsstöd 2022-1259 (OH)

The Swedish federal government under the ALF agreement 2022-Projekt0080 (OH)

## Author contributions

Conceptualization: SB

Resources: JAS

Data curation: MY

Investigation: SB, JHC, SHK, NM, VYZ, LZ, DC, OH

Formal analysis: SB, JHC, JWV

Methodology: SB, JHC, JWV

Funding acquisition: SB, NS

Supervision: AAV, LMH, NS

Writing – original draft: SB

Writing – review & editing: SB, OH, AAV, NS

## Competing interests

OH has acquired research support (for the institution) from ADx, AVID Radiopharmaceuticals, Biogen, Eli Lilly, Eisai, Fujirebio, GE Healthcare, Pfizer, and Roche. In the past 2 years, he has received consultancy/speaker fees from AC Immune, Amylyx, Alzpath, BioArctic, Biogen, Bristol Meyer Squibb, Cerveau, Eisai, Eli Lilly, Fujirebio, Merck, Novartis, Novo Nordisk, Roche, Sanofi and Siemens. AAV has received research support from BetterLife Pharma and Gilgamesh Pharma, and he has received consultancy fees from L.E.K. Consulting.

## Materials and Methods

### BV2 DKO cell line generation

Cell lines were generated from BV2 microglia cell line, a generous gift from Carol Colton (Duke university), at passage 13. Gene editing was performed using CRISPR synthetic single guide RNAs (sgRNAs, Synthego) as per manufacturer’s protocol. Briefly, cells were transfected with the Lipofectamine™ CRISPRMAX™ Cas9 Transfection Kit (Thermo) with ribonucleoprotein (RNP) complexes of synthetic sgRNAs and S. pyogenes Cas9 nuclease. RNPs were assembled with a sgRNA:Cas9 ratio of 3:3 as per the manufacturer’s suggestion. Two guides were used per gene and transfection was performed simultaneously for eIf4e-bp1 and eif4e-bp2 in a 1:1 ratio (see Table 1. for guides). After transfection, cells were incubated for 3 days in a humidified 37°C/5% CO_2_ incubator. Following incubation, cells were sorted into single cells per well in a 96-well plate. A total of 18 colonies were picked, expanded, and examined by Western blot. Double knockouts (DKO) and non-edited cells (WT) were picked as controls from the same plate and were expanded and used further for the remainder of the investigation. Mycoplasma contamination was assessed by PCR routinely.

**Table 1.**
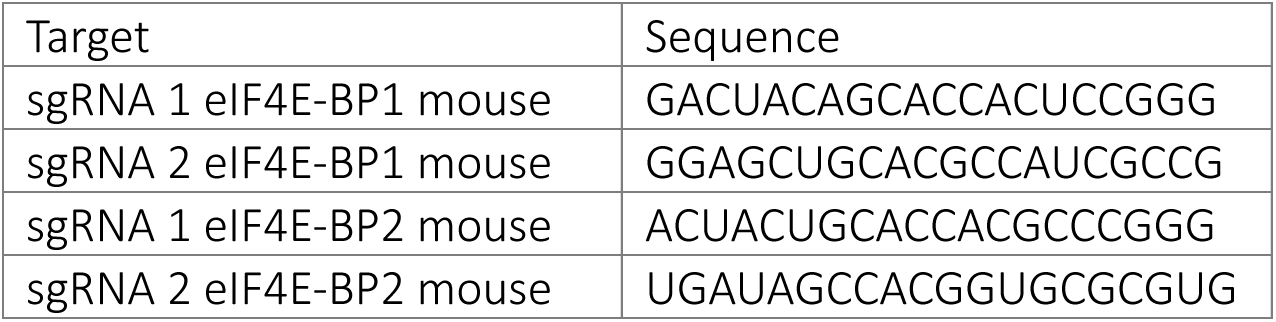
CRISPR guides used for eif4e-bp1 and eif4e-bp2 knock out in BV2 microglia cell line.

### Aβ aggregation

Aβ1–42 peptides, HFIP (A-1163-2, r-peptide) were dissolved to in anhydrous dimethyl sulfoxide (Hybri-Max D-2650 DMSO, Sigma-Aldrich) to obtain a 2.5mM Aβ stock solution, which was incubated in a bath sonicator for 10 minutes at room temperature. The peptide stock was diluted to a concentration of 100 μM in 10 mM Tris-HCl, pH 7.4, and incubated for 48 hours at 22°C to facilitate the formation of higher molecular weight oligomers, band stored at −80°C or used in experiments immediately. Individual Aβ aggregate stocks were never thawed and re-frozen. To confirm oligomeric formation, the preparation was resolved on a 4-20% TGX (Tris-Glycine eXtended) precast gel (Bio-Rad) and immunoblotted with anti-β-Amyloid 1-16 (clone 6E10; Biolegend).

### pHrodo™ Aβ labeling

For phagocytosis determination, Aβ was labeled as described by FUJIFILM Cellular Dynamics International, iCell® Microglia Application Protocol, Labeling Amyloid Beta with pHrodo Red https://www.fujifilmcdi.com/assets/iCell_MGL_ABeta_phago_AP.pdf

### In vitro treatments

BV2 cell lines were used between passages 13 and 20. Cells were maintained in Dulbecco’s Modified Eagle Medium (DMEM) (Wisent) supplemented with 10% fetal bovine serum (FBS) (Wisent) heat-inactivated (30 minutes at 60°C) and 1% streptomycin-penicillin (Wisent) in a humidified incubator containing 5% CO_2_ at 37°C. Cell culture treatments were performed in 24 or 96 well plates, at least in triplicate. BV2 microglia cells were plated at 2.5×10^4 cells/ml overnight in 10% FBS-supplemented DMEM and changed to treatment media (2% FBS DMEM) before treatment. All drugs were dissolved in treatment media, and cells were incubated with Syk inhibitor (R406, 1uM, MedChemExpress), VEH (2 uM), LPS (100ng/ml), or Aβo (2 uM) in treatment media for the indicated times. SYK inhibitor (R406, 1uM) was added 15 minutes before VEH or Aβo.

### Western blotting

Cell culture plates were washed twice with ice-cold PBS. The cells were incubated in ice-cold RIPA lysis buffer (Thermo) supplemented with an EDTA-free protease inhibitor cocktail and phosphatase inhibitor (Roche). Plates were placed in a rocking platform in the cold room for 20 minutes and scraped. After lysis, the samples were centrifuged at 12,000 g for 10 minutes at 4°C to remove insoluble material. The protein concentration was determined by the Bradford protein assay (Bio-Rad) using a BSA curve. Protein was equalized to be loaded 15ug of protein per well, with ddH_2_O mixed with an equal volume of Laemmli sample buffer. Proteins were resolved in 12 or 15% SDS-PAGE gels at 100V until the gel bottom was reached by loading buffer. Gels were transferred to 0.2 uM nitrocellulose membranes (25V overnight at 4°C), and equal protein loading was confirmed using Ponceau Red staining. The membranes were blocked for 1 hour with 5% BSA at room temperature in a rocking platform. After blocking, membranes were incubated overnight with primary antibody diluted in 5% BSA at 4°C. After 3x TBS-T (Tris-Buffered Saline, .1% Tween) washes, the membranes were incubated with peroxidase-coupled secondary antibody (1 hour at room temperature) and washed 3x on TBST. Enhanced chemiluminescence (Western Lighting® Plus ECL, PerkinElmer) was added to membranes for 1 minute (see Table 2 for antibodies). Membranes were exposed to an X-ray film.

**Table 2.** Antibodies used for Western blotting and immunofluorescence.

| Source | Antibody | Catalogue # |
| --- | --- | --- |
| CST | 4E-BP1 | #9644 |
| CST | Phospho-4E-BP1 (Thr37/46) | #2855 |
| CST | Phospho-4E-BP1 (Ser65) | #9451 |
| CST | 4E-BP2 | #2845 |
| CST | Phospho-Syk (Tyr525/526) | #2710 |
| CST | Syk | #2712 |
| CST | NF-κB p65 | #8242 |
| R&D Systems | IL-1b | AF-401-NA |
| Sigma | b-Actin | #A5441 |

### Live-Cell Incucyte imaging

Microglia were plated into a 96-well plate at 2.5×10^4 cells/ml (4 to 5 wells per cell line/condition). At time 0, wells were treated with Caspase-3/7 (CellEvent™ Caspase-3/7 Green, Invitrogen) 1:1000 in treatment media, as per manufacturer instructions, with Aβ, pHrodo™ labeled Aβ or VEH. Four 20X images per well were collected every hour. Using Incucyte® S3 Live-Cell Analysis System software, 2019A image masks for phase confluence, caspase 3/7 signal (green), and pHrodo™ labeled Aβ (red) were generated. Graphs display caspase normalized to phase confluence.

### Immunocytochemistry

Cells were seeded on poly-L-lysine (Sigma) coated coverslips. After the indicated treatments, cells were washed in PBS and fixed in 4% paraformaldehyde (PFA) for 15 minutes on a rocking platform at room temperature. After washing 3X with PBS, cells were permeabilized with PBS containing 0.05% Triton X-100 for 15 minutes at room temperature and blocked with blocking solution (PBS containing 10% BSA). Fixed cells were incubated with the primary antibody in a blocking solution overnight at 4°C and thereafter incubated with Alexa Fluor (Thermo)-conjugated secondary antibodies (1:1000 in blocking solution) for 1 h at room temperature protected from light (see Table 2 for antibodies). Coverslips were mounted onto a glass slide using the ProLong Diamond Antifade Mountant with DAPI (Thermo). For Mitotracker green (Thermo) staining, live imaging was used, and no fixing was performed. Mitotracker Green was dissolved into a final concentration of 100 nM in phenol red-free FBS-free DMEM medium according to the manufacturer’s instructions, and cells were incubated for 30 min at 37°C. Samples were imaged with a ZEISS confocal microscope (LSM 800). Image processing was performed with FIJI (NIH).

### Cytokine measurement

Media was obtained from treated cells as indicated, centrifuged at 10,000 rpm for 5 minutes at 4°C, aliquoted, and frozen. Samples were shipped in dry ice to Eve Technologies Corporation (Calgary, Canada). The cytokines GM-CSF, IFNγ, IL-1β, IL-2, IL-4, IL-6, IL-10, IL-12p70, MCP-1 and TNFα were evaluated in a Mouse Cytokine Pro-Inflammatory Focused 10-Plex Discovery Assay® Array (MDF10). Cytokines with less than 3 readings/group or values below detection were excluded from the results.

### Lactate measurement

Cells were plated in 96-well cell culture plates at 2.5×10^4 cells/ml (4 to 5 wells per cell line/condition) and treated as indicated with Aβo or VEH. Lactate secreted into the cultured medium was quantified using a Lactate Lactate-Glo™ (Promega) according to the manufacturer’s instructions.

### Measurement of Real-Time OCR

Real-time oxygen consumption rate (OCR) was measured using XFe96 plate (Seahorse Bioscience) according to the manufacturer’s instructions, with minor modifications. Briefly, cells were plated on a XFe96 cell culture microplate and treated for 4 hours with Aβ or VEH. The cartridge plate was hydrated with XF calibrant buffer and incubated overnight (37°C, CO_2_-free). The assay medium (XF base medium containing 1 mM pyruvate, 4 mM glutamine, and 25 mM glucose) was prepared immediately before the assay. XF sensor cartridges were loaded with test compounds. OCR was measured every 6 minutes in response to the ATPase inhibitor oligomycin (2.5 uM), the uncoupling agent FCCP (1 uM), and the electron-transport-chain inhibitors rotenone (2 uM), and Antimycin A (1 uM).

### Animals and environment

Mice were housed in standard laboratory cages with 4–5 mice in each cage. Mice were given water and standard rodent chow ad libitum. Cages were maintained in ventilated racks in temperature-(20–21 °C) and humidity-(∼50%) controlled rooms, on a 12 h light/dark cycle. Standard corncob bedding was used for housing (Harlan Laboratories, Inc). Mice were maintained under standard conditions at the Goodman Cancer Institute (GCI) animal facility. All procedures followed the Canadian Council on Animal Care guidelines and were approved by the McGill University Animal Care Committee.

### Mouse lines

Eif4ebp1^f/f^ and Eif4ebp2^f/f^ mice are an established mouse line in the laboratory (Perth, Australia)(25). Mice were generated by Ozgene Pty Ltd (Perth, Australia) by targeting a construct containing a neomycin selection cassette flanked by two short flippase recognition target (FRT) sites was used to generate a conditional allele (25). The targeting construct also contained two loxP sites between exons 1 and 2 and between exons 2 and 3 for both Eif4ebp1 and Eif4ebp2 genes. The neomycin selection cassette was removed after the generation of the ‘floxed’ lines. Conditional microglial knockouts were generated by crossing Eif4ebp1^f/f^ Eif4ebp2^f/f^ with the B6.129P2(Cg)-Cx3cr1tm2.1(cre/ERT2) Litt/WganJ mouse line (Cx3cr1^creERT2^, Jackson laboratories) and then further crossed with (JAX stock # 011029 B6N.129-Rpl22^tm1.1Psam^/J, Jackson laboratories). The presented translatomic and transcriptomic data was generated with mice homozygous for the Rpl22^HA^ allele and heterozygous for the Cx3cr1^creERT2^ allele.

### Genotyping

During weaning, ear punches were collected from 3-week-old pups to determine the genotype. The DNA was prepared by boiling tissue in 50 uL of Solution 1 (25mM NaOH, 0.2mM EDTA, pH 12.0) for 20 minutes at 100°C. Next, 50uL of Solution 2 (40mM HCl, pH 5.0) was added and the tube was vortexed. The PCR reactions were performed using AccuStart™ II PCR SuperMix (Avantor) following the manufacturer’s instructions (see Table 3 for primers). The reactions were carried out and the PCR products were visualized on a 1.5% TAE-Agarose gel with ethidium bromide.

**Table 3.**
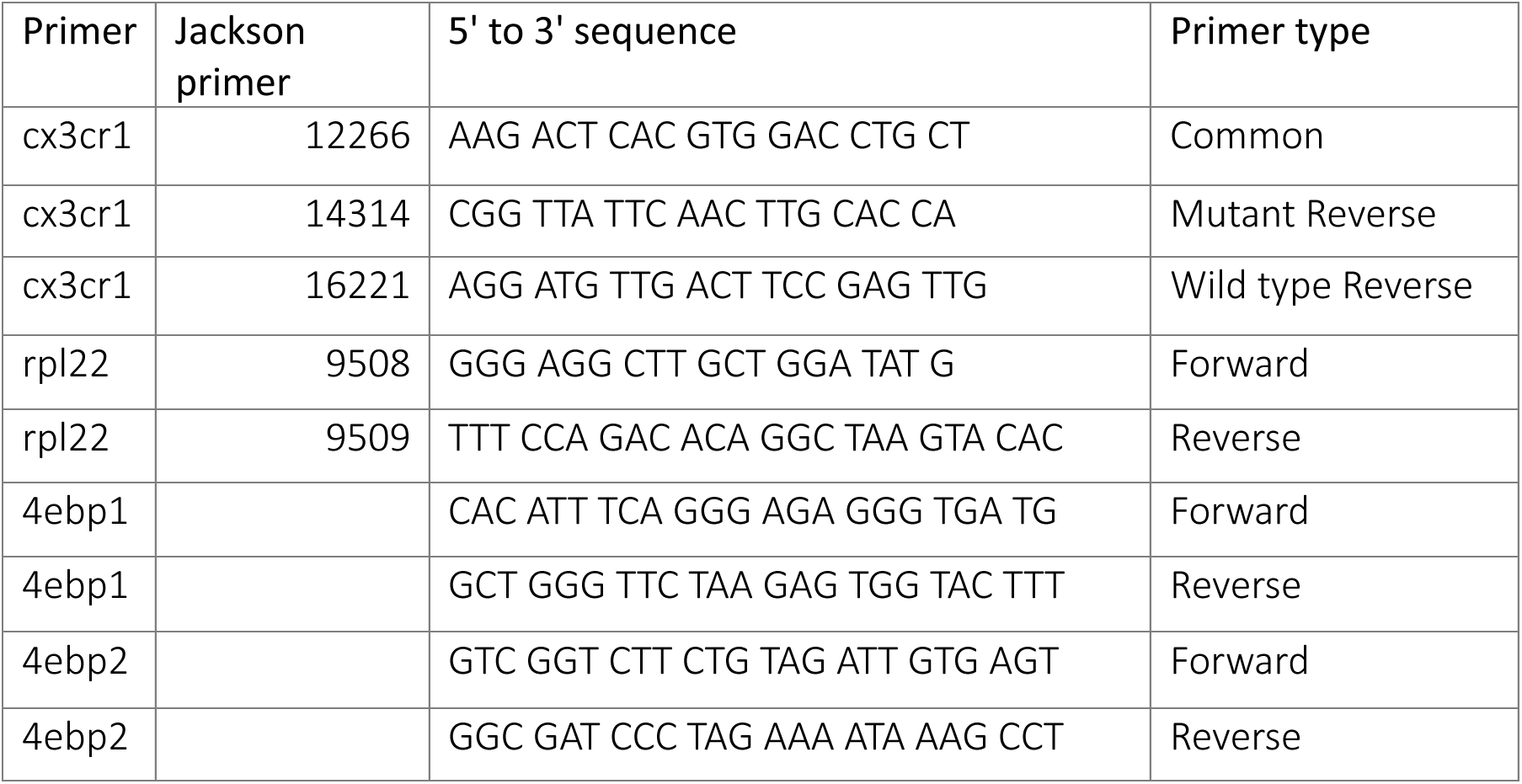
Primers used for mouse model genotyping.

### Immunohistochemistry

Mice were placed under isoflurane anesthetics until loss of pain reflex and transcardially perfused with filtered ice-cold PBS then 4% paraformaldehyde (PFA, Electron Microscopy Sciences). Brains were cryoprotected in 30% sucrose. Brains were sectioned coronally into 30 μm-thick slices on a freezing microtome (Leica SM 2010R) and stored in a solution of 0.05% NaN3 PBS as free-floating slices. For immunostaining, tissue was blocked for 1 h in PBS, 0.2% Triton X-100, and 10% goat serum. Immediately following blocking, brain sections were incubated with primary antibodies diluted in PBS supplemented with 1% goat serum and incubated overnight at 4 °C. Samples were then incubated with Alexa Fluor (Thermo)-conjugated secondary antibodies (1:1000 in blocking solution). Sections were adhered to glass slides and coverslips were mounted onto a glass slide using DAPI Mounting Medium (VECTASHIELD). Samples were imaged with a ZEISS confocal microscope (LSM 800). Image processing was performed with FIJI (NIH).

### Ribosome pulldown

Bilateral hippocampi were extracted from mice, flash-frozen in liquid nitrogen, and stored at − 80°C until use. Simultaneously, samples were homogenized in ice-cold homogenization buffer (50 mM Tris, pH 7.4, 100 mM KCl, 12 mM MgCl2, 1% NP-40, 1 mM DTT, 1:100 protease inhibitor, 200 units/ml RiboLock RNase Inhibitor (Thermo) and 0.1 mg/ml cycloheximide in nuclease-free H2O (Wisent) 10% w/v with a dounce homogenizer (Sigma) until a homogeneous suspension could be observed. The sample was incubated for 10 minutes on ice, and 1 ul of the lysate was transferred to a microcentrifuge tube and centrifuged at 10,000 g at 4°C for 10 minutes. Supernatants were transferred to a fresh microcentrifuge tube on ice, and then 40ul was removed for input fraction analysis. Two ug of IgG (Sigma, only for validation pilot) or anti-HA (Biolegend) were added into the lysate and incubated for at least 30 minutes at 4°C in a rotator. Meanwhile, Dynabeads (Thermo), 100 ml per sample, were equilibrated to homogenization buffer by washing 3X. At the end of the sample incubation with antibody, beads were added to each sample, followed by incubation overnight at 4°C on a rotating mixer. The next day, samples were washed 3X with high-salt buffer (50 mM Tris-HCl, pH 7.4, 300 mM KCl, 12 mM MgCl2, 1% NP-40, 1 mM DTT, 1:200 protease inhibitor, 100 units/ml RiboLock, and 0.1 mg/ml cycloheximide in nuclease-free H2O (Wisent) and magnetized. Beads were magnetized and immobilized, and excess buffer was removed and 700μl of Trizol was added. RNA was eluted using the Direct-zol RNA Microprep kit (Zymo Research) per manufacturer’s instructions.

### RNA Sequencing

RNA-Seq was performed at the University of Toronto Donnelly Centre. Two or three samples were pooled, and libraries were constructed using the NEBNext Ultra II Directional RNA Library Prep Kit for Illumina. Sequencing was performed on an Illumina NovaSeq 6000 system for a total of 24 samples (n=3-4/group, IP). Canadian Center for Computational Genomic’s pipeline GenPipes (58) was used to align the raw files and quantify the read counts. Briefly, raw fastq files were aligned to the mouse genome GRCm38 Genome Reference with default parameters and raw reads were quantified using HTseq count (59).

### Differentially expressed genes analysis

The pipeline for differentially expressed genes is as follows. First, genes with low read counts (<10) were filtered out, which brought the total number of genes from 47,069 to 24,557. Next, we took two steps for visual quality control of the samples. By inspecting the log2-normed sample by gene heatmap, one sample (DKO) showed severely outlying values across all genes and was excluded. As a second quality control step, we performed cell-type deconvolution of each sample to assess whether IP samples showed microglia gene enrichment. We used the ‘cellanneal’ python package (60), while using the Allen Mouse Brain Atlas “Whole cortex & hippocampus with 10x-smart-seq taxonomy” atlas as the reference dataset (50), samples showing aberrant deconvolution indicated compromised sample quality. The 3 samples identified through these two quality control procedures were removed.

The “pyDESeq2” python library was used for differential expression analysis. Briefly, this process involves fitting size factors, dispersions, and dispersion trend curves to identify log-fold change across given contrasts. The model was fit once to identify outliers and then once again after outliers had been refit. We evaluated a WT vs. DKO contrast. We report 88 genes surviving multiple comparisons correction (p[adj]<0.05).

### Human cerebrospinal fluid biomarker analysis

To investigate whether our findings generalize to human Alzheimer’s disease, we examined associations between biomarkers measured in the CSF of 688 healthy and AD-spectrum individuals from the Swedish BioFINDER-2 cohort (NCT03174938)(61). Individuals had a clinical diagnosis of either cognitively normal (CN), subjective cognitive decline (SCD), mild cognitive impairment (MCI), or Alzheimer’s dementia (AD). All were recruited from the Skåne University Hospital and the Hospital of Ängelholm, Sweden. Participants were included in the study if they had both OLINK and neurotoolkit CSF biomarker data available. MCI and AD patients were only included if they showed biomarker evidence for Alzheimer’s disease, namely CSF Aβ42/40 ratio below the previously established abnormality threshold of 0.08 (62). Detailed inclusion criteria for the BioFINDER-2 study were described elsewhere (63). CSF was acquired through lumbar puncture. Biomarkers for Aβ40, Aβ42, neurofilament light (NfL) and sTREM2 were measured as part of the NeuroToolKit assay panel in accordance with the manufacturer’s instructions (Roche Diagnostics International Ltd)(64). Technicians collecting and processing data were blind to clinical and biomarker information. CSF EIF4EBP1 was measured using proximity extension assay as part of the OLINK Explore 3072 panel, developed by OLINK Proteomics (Uppsala, Sweden) EIF4EBP1 expression was log2 transformed and z-transformed. Demographic information and CSF levels stratified by clinical diagnosis can be found in Table 4.

**Table 4.** Demographic characteristics of included study participants. Cognitively normal (CN), Subjective Cognitive Decline (SCD), Mild Cognitive Impairment (MCI), Alzheimer’s Dementia (AD)

|  | CN | SCD | MCI | AD | Total |
| --- | --- | --- | --- | --- | --- |
| N | 256 | 130 | 140 | 162 | 688 |
| Age (SD) | 62.8 (16.1) | 67.7 (8.7) | 72.2 (7.6) | 73.4 (7.1) | 68.2 (12.4) |
| Females (%) | 147 (57.4%) | 59 (45.3%) | 69 (49.2%) | 87 (53.7) | 362 (52.6%) |
| Abnormal Aβ42/40 Ratio (%) | 48 (18.8%) | 69 (53.1%) | 140 (100%) | 162 (100%) | 419 (60.9%) |
| APOE e4 Carrier (%) | 99 (38.7%) | 69 (53.1%) | 101 (72.1%) | 112 (69.1%) | 381 (55.4) |
| CSF NFL (SD) | 127.0 (95.5) | 172.7 (108.8) | 228.3 (139.5) | 325.2 (214.9) | 200.4 (161.4) |
| CSF sTREM2 (SD) | 10.3 (3.3) | 11.2 (3.1) | 11.7 (3.8) | 13.1 (4.1) | 11.4 (3.7) |
| z CSF EIF4EBP (SD) | -0.1 (0.4) | 0.0 (0.5) | 0.0 (0.4) | 0.1 (0.5) | 0.0 (0.5) |
| CSF Aβ40 (SD) | 19.6 (5.3) | 20.2 (5.9) | 20,1 | 19,4 | 19.8 (5.9) |

General linear models assessed the interaction between CSF EIF4EBP1 and sTREM2 levels on CSF NfL levels, including for age and sex as covariates of no interest. Individual variation in CSF levels can be driven by non-disease related physiological differences unless adjusted using a “reference protein” (65). Therefore, the model additionally included Aβ40 as a covariate of no interest, as this protein was found to be an appropriate reference protein across several datasets. The level for a significant interaction was set 0.05. To interpretation and visualization only, participants were divided into one of three equal-sized CSF EIF4EBP1 categories (high, medium, low) based on a quantile cut of ranked values. However, in the model, EIF4EBP1 was expressed as a continuous variable.

### Quantification and statistical analysis

Statistical analyses were performed using GraphPad Prism 9.0 (GraphPad Software). Comparisons between two groups were performed using two-tailed unpaired t-test. One-way analysis of variance (ANOVA) with Tukey’s post hoc test was used to compare three or more independent groups. For comparison of multiple factors, a two-way ANOVA followed by Tukey’s post hoc test was used. Data are presented as mean ± standard error of the mean (SEM). Statistical parameters are detailed in the legend for each figure.

