## Supplementary figures for "Translational control of microglial inflammatory and neurodegenerative responses"

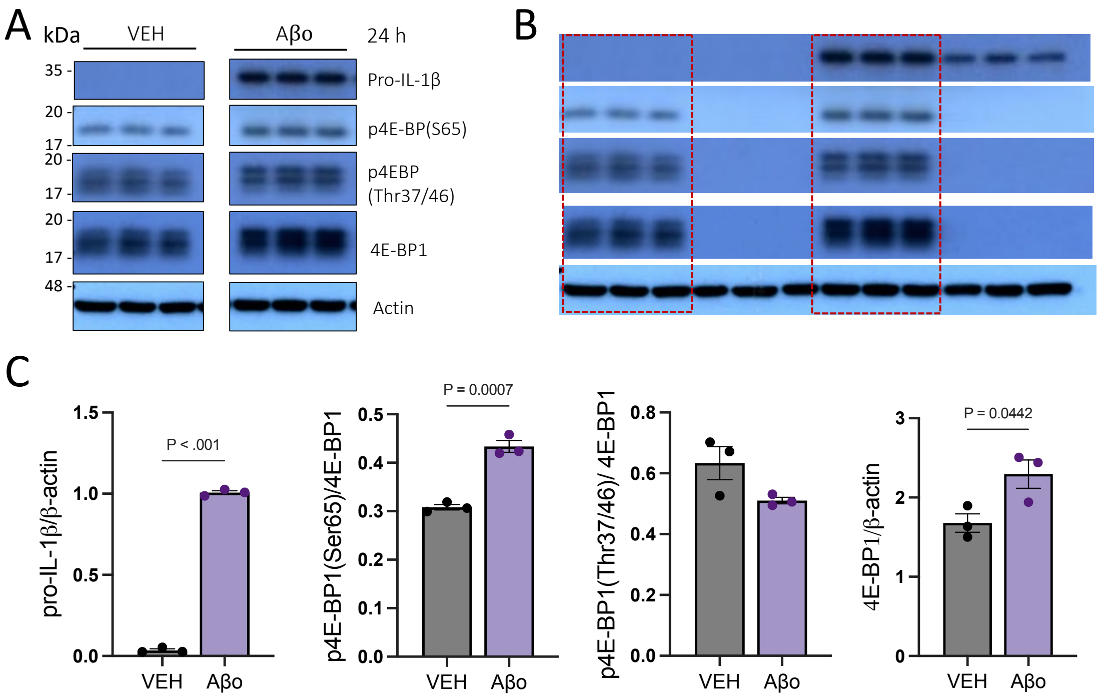


Fig. S1 Prolonged Aβo exposure triggers 4E-BP1 expression. (A) Representative western blot analysis of total and phosphorylation status 4E-BP1, and β-actin in BV2 cell lysates 24h after vehicle (control) or Aβo (2 μM) treatment. (B) Uncropped Western blot. (C)Quantification of phosphorylation ratio to total protein or to β-actin loading control (n=3/group). Data are presented as means ± SEM (P values were calculated using two-tailed unpaired t-test).

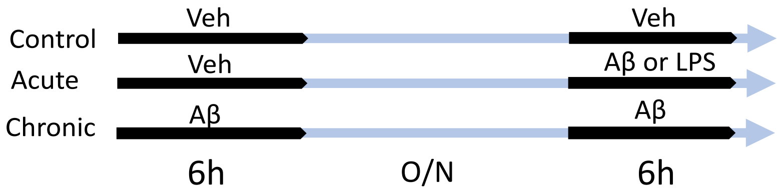


Fig. S2 experimental design for the chronic model study. Experimental design used for the chronic model study (modified from Baik, S.H., et al., 2019).

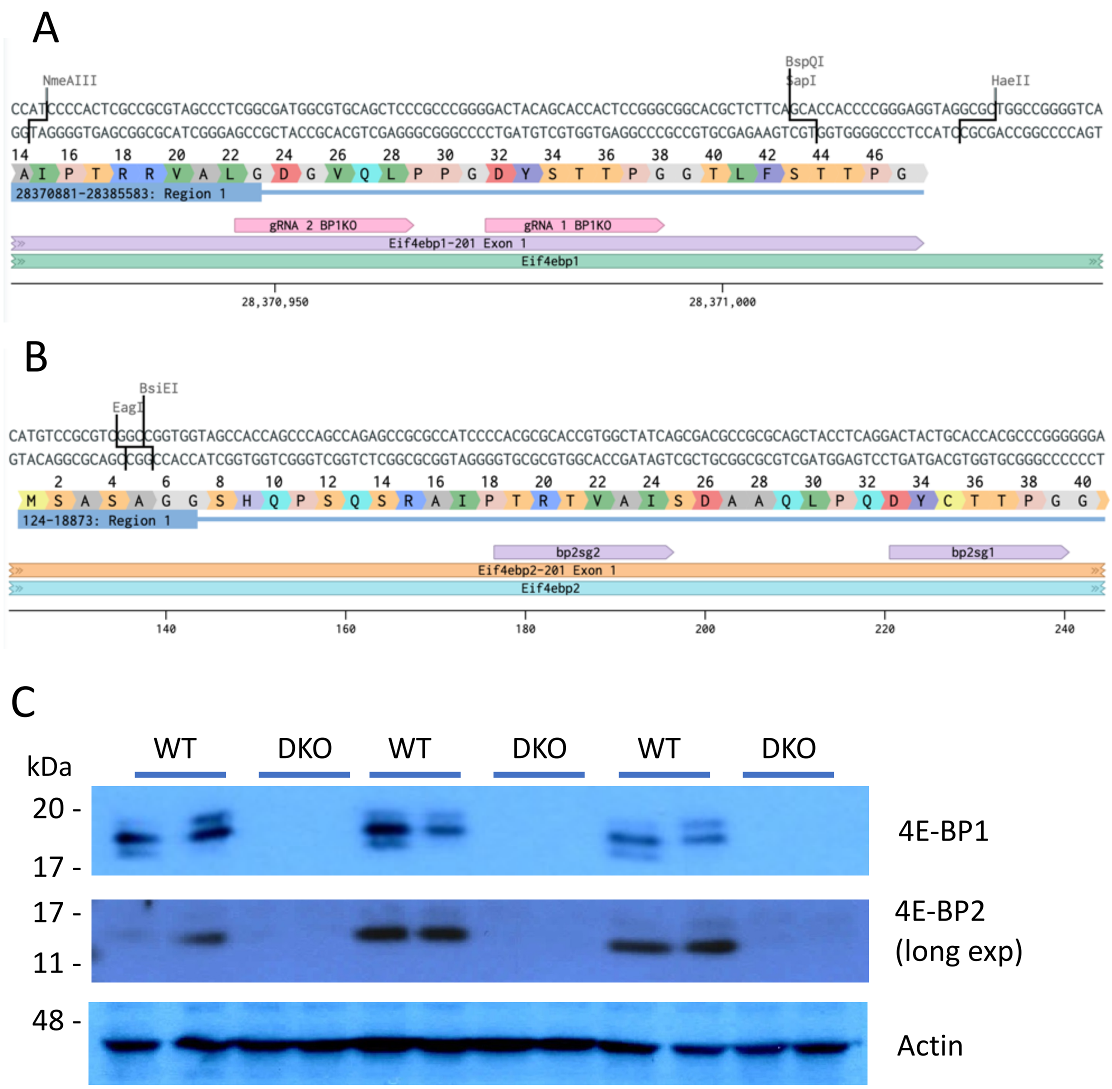


Fig. S3 Generation of Eif4ebp1 and Eif4ebp2 cell lines. a and b, Schematic representation of the targeting guides used for the generation of Eif4ebp1 and Eif4ebp2 knockout in BV2 microglia. c Knock out confirmation of selected colonies.

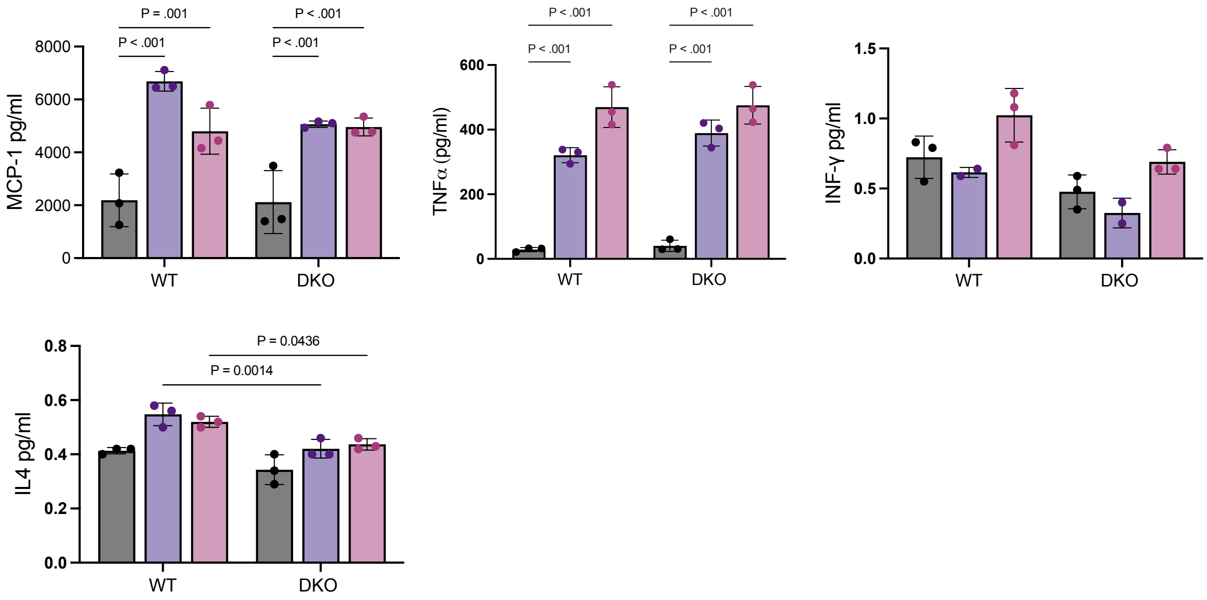


Fig. S4 Cytokine excretion examination. Determination of cytokines in supernatant from Aβo (24h) or LPS (6h) treated microglia. Data are presented as means ± SEM (P values were obtained using two-way ANOVA, Tukey's post-hoc test).

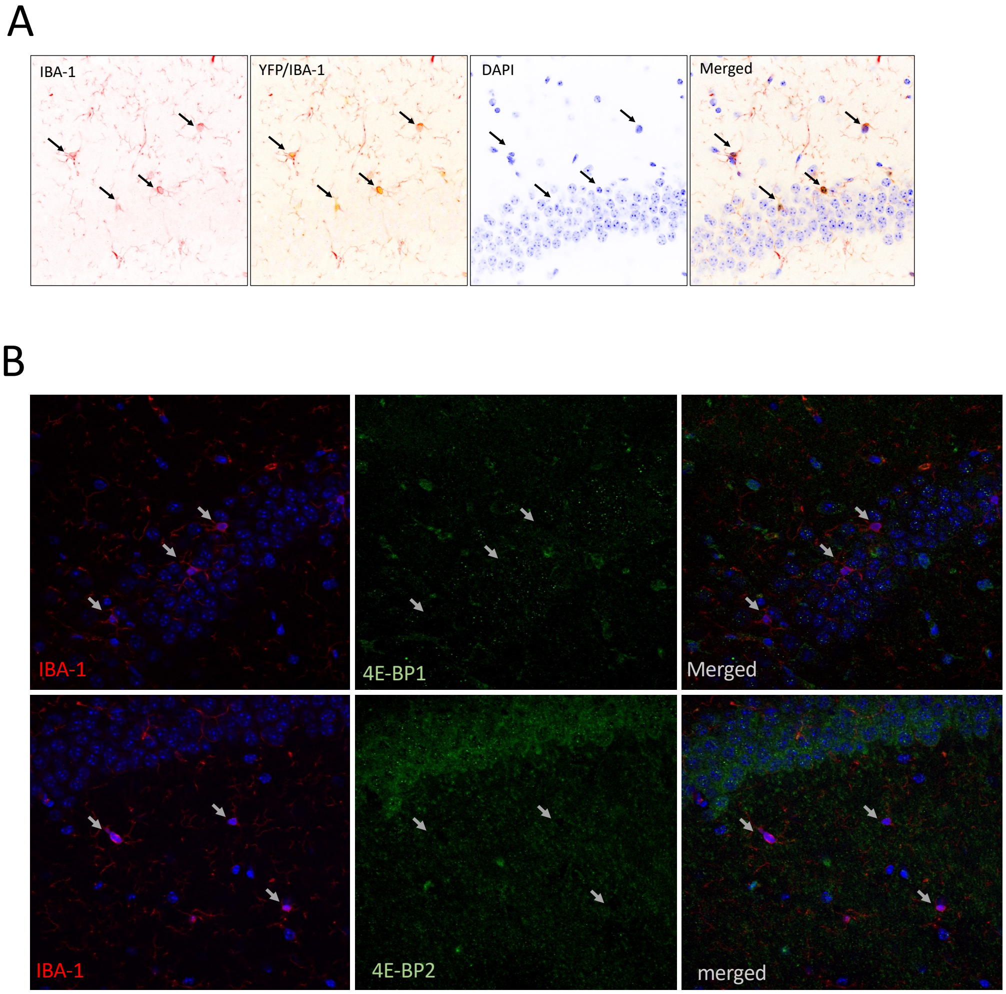


Fig. S5 In-vivo model Knock out validation. (A) Representative confocal microscopy images of Cx3cr1$^{CreER/+}$ mouse expressing YFP as marker of CRE recombinase expression in microglia. (B) Representative confocal microscopy images of DKO microglia in Cx3cr1$^{CreER/+}$; 4E-BP1$^{f/f}$/4E-BP2$^{f/f}$ mice 2 weeks after tamoxifen induction.

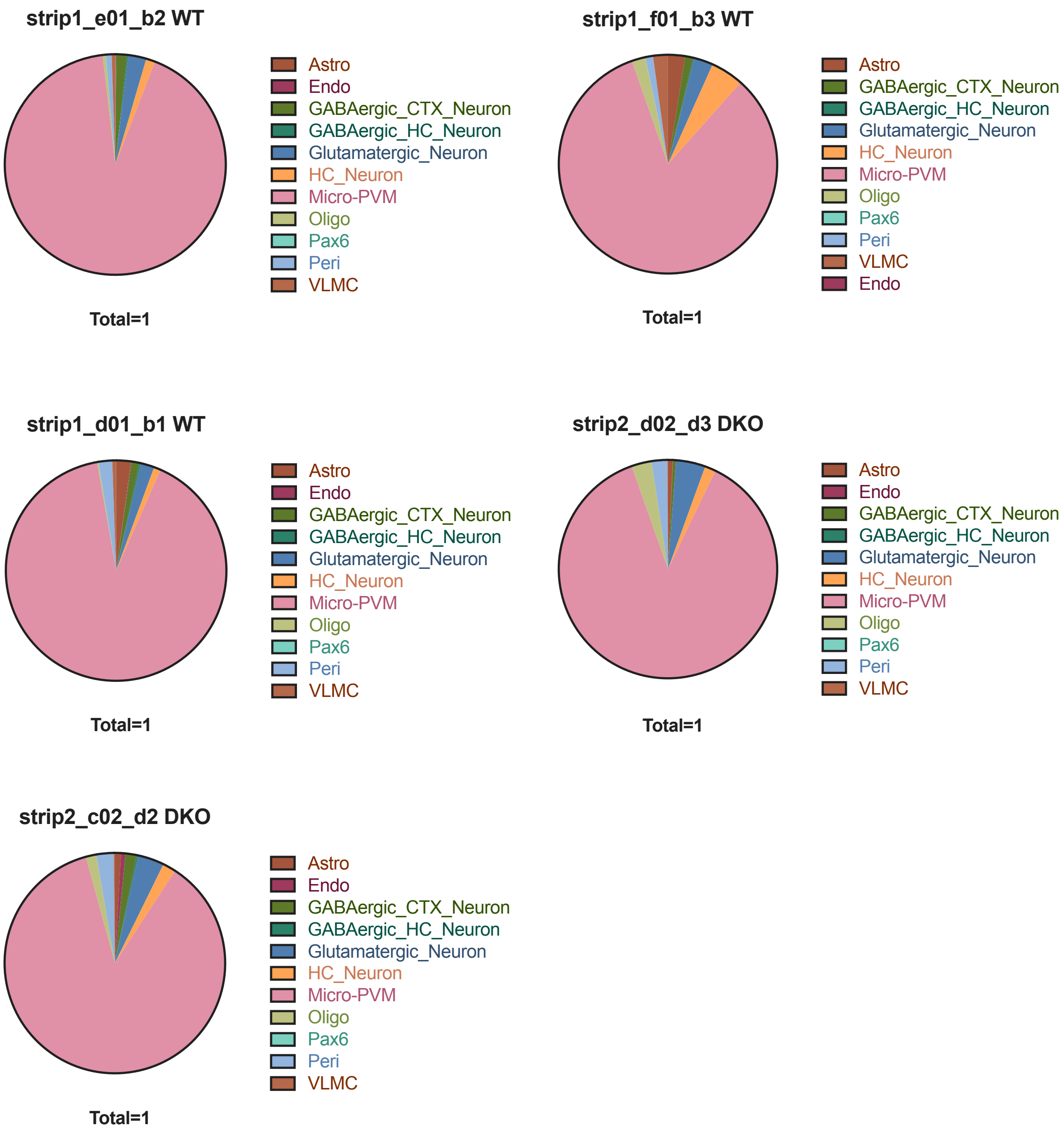


Fig. S6 Cell-type deconvolution. Cell type enrichment pie chart per sample.

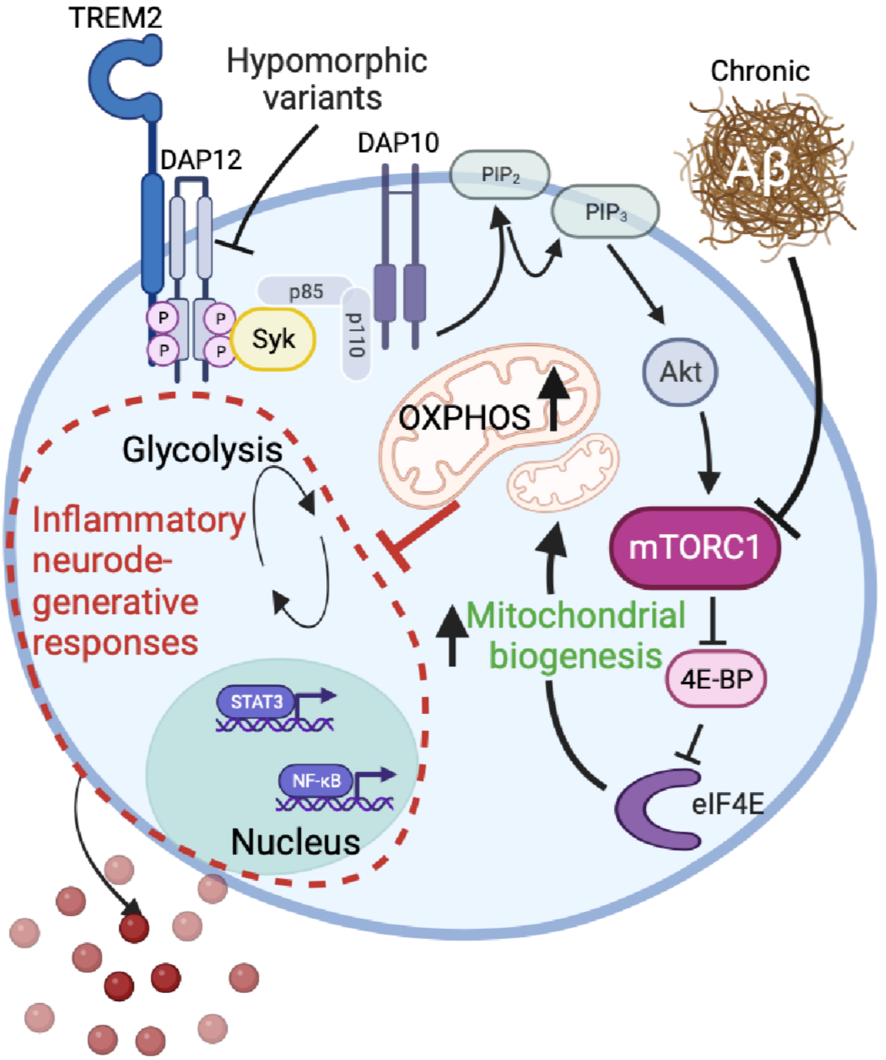


Fig. S7 proposed model for mTORC1-4E-BP1 axis role in microglia. mTORC1-mediated inactivation of 4E-BP1 limits microglia pro-inflammatory output in response to Aβ by stimulating mitochondria biogenesis and pivoting the associated glycolytic energy pathway towards OXPHOS-dependent metabolism. mTORC1-mediated inactivation of 4E-BP1 acts as a mechanism of immune resolution and, ultimately, neuroprotection. This mechanism is dependent on TREM2/SYK signaling and decreases during chronic exposure to Aβo. Thus; we propose the engagement mTORC1-4E-BP1 axis as a potential therapeutic target for microglia modulation in AD. In addition, 4E-BP1 may provide a useful biomarker to monitor microglia state and modulation. Created with BioRender.com.
